# Mutation of the allexivirus-specific P40 gene of Wineberry latent virus, an isolate of Blackberry virus E, greatly reduces virus multiplication

**DOI:** 10.1101/2024.06.18.599508

**Authors:** Wendy McGavin, Graham Cowan, Susan Jones, Stuart MacFarlane

## Abstract

The complete sequence of Wineberry latent virus (WLV), a previously reported but uncharacterised *Rubus*-infecting virus with flexuous particles, has been determined. Analysis shows WLV to have 76% overall nucleotide sequence identity to the more recently discovered Blackberry virus E, an allexivirus belonging to the subgroup of these viruses that lack the 3’ proximal cysteine-rich protein (CRP) gene present in *Allium*-infecting allexiviruses. An infectious cDNA clone of WLV was constructed and a mutation introduced into the P40 gene, which is an allexivirus-specific gene of unknown function. In addition, the infectious clone was modified to express the green fluorescent protein (GFP) as an N-terminal fusion to the WLV coat protein (CP). Using this GFP overcoat strategy it was possible to follow the multiplication and movement of the virus in infected *Chenopodium quinoa* and spinach leaves. Introduction of the frameshift mutation into the P40 gene of WLV reduced virus accumulation by 97%, and with the GFP overcoated WLV the P40 mutation almost entirely abolished GFP fluorescence in inoculated leaves suggesting that the WLV P40 protein is required for normal levels of virus multiplication.

**Repositories:** WLV complete sequences deposited at GenBank. Accession Nos. MZ944847.1, OQ877124.1

## Introduction

Wineberry latent virus (WLV) was first reported in Scotland in one plant growing at the James Hutton Institute among a group of Japanese wineberry (*Rubus phoenicolasius*) plants that had been imported from the US more than 10 years previously (Jones, 1977). The virus was symptomless in wineberry but produced necrotic local lesions when mechanically inoculated to *Chenopodium quinoa*, with symptomless systemic infection developing occasionally. Systemic infection occurred after mechanical inoculation to three other *Chenopodium* species and local infection only occurred in five of nineteen other herbaceous species that were tested. Mechanical transmission to red raspberry (*R. idaeus*), black raspberry (*R. occidentalis*), wineberry and *R. procerus* was not successful, however, transmission by grafting to some red raspberry varieties, to *R. occidentalis, R. loganobaccus, R. procerus* and to wineberry was successful. In all of these cases no symptoms of infection were seen in the *Rubus* plants and WLV was detected only by back-inoculation to *Ch. quinoa*. In a later study, WLV was found to have flexuous filamentous particles of c. 620 × 12 nm that did not react to a panel of antibodies raised against more than twenty other filamentous viruses (Jones *et al*., 1990).

Raspberry and blackberry plants are produced by clonal propagation for commercial fruit production and breeding purposes and internationally agreed pathogen testing procedures are in place to prevent the release of diseased plants into the production system. Since its discovery, WLV has remained listed as a pathogen of *Rubus* plants and is tested for by a mechanical inoculation to indicator plants such as *Ch. quinoa*, which is expected to detect a wide range of mechanically-transmissible viruses. The initial aim of the current study was to determine the sequence of the WLV genome and so confirm its identity and make a molecular diagnostic test possible. Thereafter an infectious clone of WLV was constructed, the first example of a systemically infectious cDNA clone of an allexivirus from outside the allium-infecting sub-group, enabling a mutational study of the role of the WLV-encoded allexivirus-specific P40 gene in virus infection.

## Methods

### Recovery and multiplication of WLV

A freeze-dried sample of WLV-infected *Ch. quinoa* leaves, collected in 1973, was homogenised in water and mechanically inoculated to *Ch. quinoa* and *Ch. amaranticolor* plants, producing necrotic lesions (Fig 1.). This infection was further passaged to multiply the virus, and virus particles were isolated from the infected leaves as described previously (Jones *et al*., 1990).

**Figure 1.**
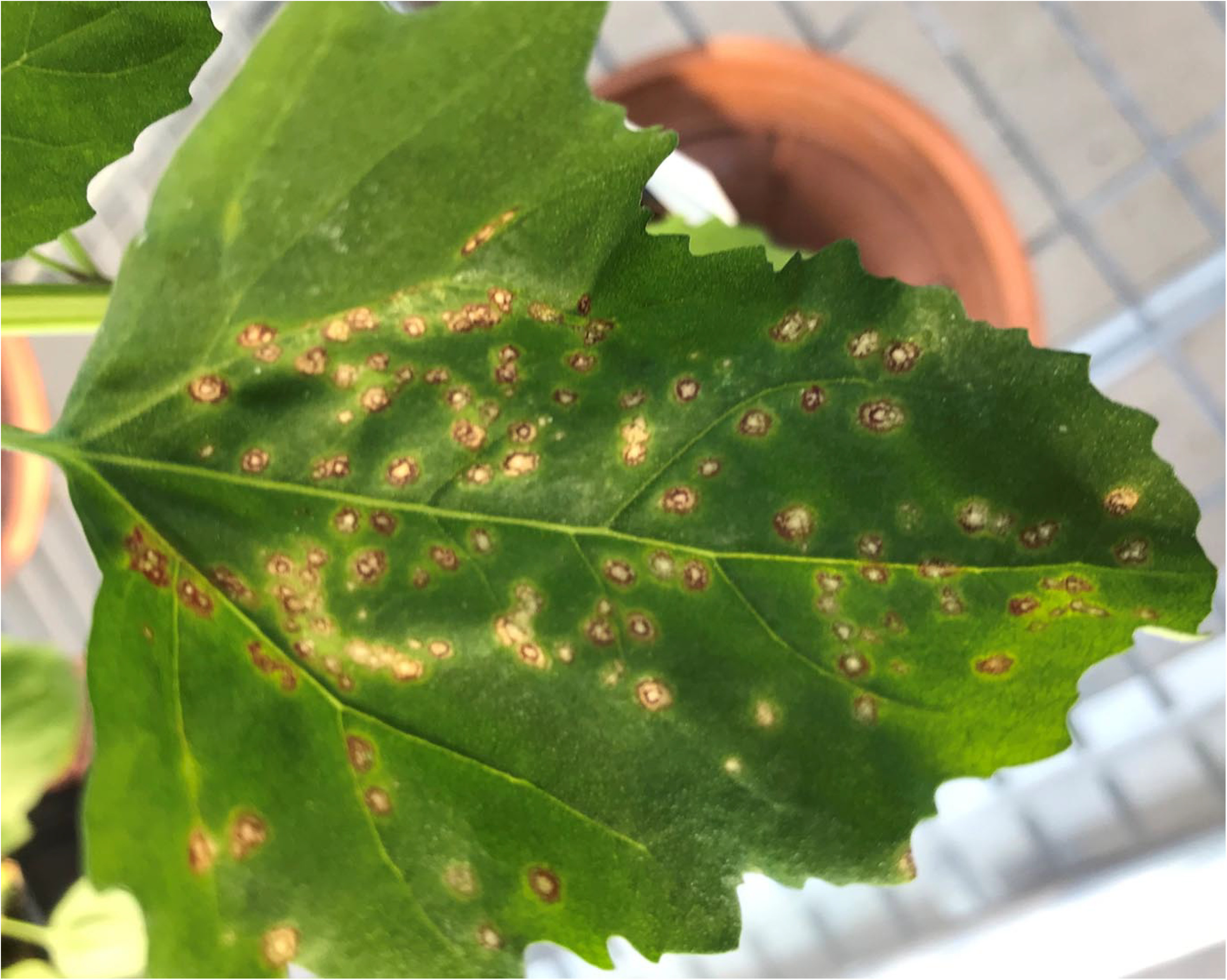
Necrotic lesions caused by WLV in *Ch. amaranticolor* at 13 days post inoculation.

### Sequencing of WLV

Viral RNA was isolated from the purified virus particles using a SDS and phenol extraction followed by ethanol precipitation. Double-stranded cDNA was prepared from this RNA using SuperScript™ IV Reverse Transcriptase (Invitrogen), a first strand random hexamer primer linked to an Oxford Nanopore Technologies (ONT) adaptor sequence (PCR-RH-RT) and a strand-switching ONT primer (PCR-Sw_mod_3G) to initiate second strand synthesis as described by Young *et al*. (2019) (Supplementary Table 1).

The cDNA was amplified to incorporate barcoding sequences using the ONT PCR Barcoding kit (SQK-PBK004) and recommended protocol. The amplified cDNA was loaded onto a ONT Flongle device (24hr read). Base calling was conducted using Guppy (Oxford Nanopore Technologies, 2019), a quality assessment was conducted using r-minionqc (Lanfear *et al*., 2019) and reads were trimmed using NanoFilt (De Coster *et al*., 2018) with the parameters: -q 10 -l 500 --headcrop 50. The trimmed reads were then aligned against a dataset of 1,431,042 proteins derived from 31,357 viruses with complete genomes in the virus partition of the NCBI’s genome assembly database (based on v240 of Genbank) using DIAMOND (v0.9.35). DIAMOND was used with parameters designed for mapping long DNA sequences that may span several genes (blastx max-target-seqs 5 e-value 0.000001, -F 15, range-culling, top 10, subject-cover 25, sensitive).

For confirmation of the ONT-derived sequence, the complete genome of WLV was amplified using Phusion DNA polymerase (New England Biolabs) from total RNA of WLV-infected *Ch. quinoa* purified using the Nucleospin plant RNA kit as directed by the manufacturer (Machery-Nagel). Amplification primers for the cloning and sequencing were designed from selected ONT sequences and from re-sequencing of amplified fragments. The 3’ terminus of WLV RNA was amplified from cDNA synthesised using an oligo-dT primer. The 5’ terminus of the WLV RNA was obtained by G-tailing of full-length first strand cDNA using terminal transferase and amplification using an oligo-dC primer combined with a virus-specific minus-strand primer, as described previously (McGavin and MacFarlane, 2008).

### Construction of a full-length cDNA clone of WLV

A 122 nt DNA fragment consisting of 21 nt from the 3’ end of the CaMV 35S promoter linked to 23 nt from the 3’ terminus of WLV followed by a 49 nt polyA sequence and 29 nt from the HDV ribozyme was created by PCR amplification of a mixture of 3 primers (Supplementary Table 1): primer 4347 (35S+3’WLV), primer 4104 (HDV ribozyme, minus strand), primer 4344 (3’ end WLV+polyA49+HDV ribozyme). This fragment was inserted by assembly reaction (NEBuilder HiFi DNA Assembly Mix, New England Biolabs) between the CaMV 35S promoter and HDV ribozyme sequences of the small, low copy number binary plasmid pDIVA (Laufer et al., 2018) that had been amplified by inverse PCR using primers 2740 (minus strand CaMV 35S promoter) and 2757 (plus strand HDV ribozyme) to create intermediate plasmid p2073. The complete genome of WLV was inserted into p2073 by several rounds of assembly reaction using four separate overlapping PCR amplified WLV fragments to produce several clones of the full-length pDIVA/WLV plasmid p2080. During the sequential inverse PCR of the vector plasmid the polyA sequence was reduced (presumably by polymerase slippage) to 42 nt. The full-length WLV clone (referred to as p2080 v5.9) was fully sequenced and transformed into *Agrobacterium tumefaciens* strain AGL1 for plant infection.

### Construction of GFP overcoated WLV and mutants

The c. 3.2 kb 3’ terminal region of WLV from p2080 is flanked by unique NsiI and SalI restriction sites and this DNA fragment was used for insertion of GFP and gene mutation. The GFP gene with an in-frame foot and mouth disease virus (FMDV) 2A autocatalytic peptidase domain at its 3’ end was PCR amplified from pTXS.GFP-CP (Santa Cruz et al., 1996), using primers 4456 (WLV CP promoter+5’GFP plus sense) and 4458 (FMDV 2A+WLV CP minus sense) (Supplementary Table 1) and inserted by assembly reaction upstream of and in-frame with the second (proline) codon of the WLV CP gene. This creates a 16 amino acid FMDV 2A peptide linker (NFDLLKLAGDVESNPG) between the GFP protein and the WLV CP. The entire WLV 3’terminal region carrying the GFP-2A insertion was sequenced before being re-cloned into p2080 to create p2108.

A frameshift mutation was introduced into the P40 gene of the wild-type full-length WLV clone p2080 and also the GFP overcoat WLV clone p2108 by digestion at a BsrGI restriction site, infilling using T4 DNA polymerase and plasmid religation to produce the mutant clones p2106 and p2111, respectively. In addition, a frameshift mutation was introduced at a MluI restriction site in the ORF1 (replicase gene) of clone p2080 to produce clone p2107. This mutation was expected not to affect transcription of WLV RNA from the agroinfiltrated binary plasmid but to prevent replication of WLV.

### Inoculation of plants by agroinfiltration and bombardment

48 hr cultures of *A. tumefaciens* AGL1 carrying the pDIVA/WLV plasmids were resuspended in infiltration buffer (10 mM MgCl2, 10 mM MES pH 5.7, 150 µM acetosyringone) at an OD_600_ = 0.2, incubated at room temperature (c. 25°C) for 2-3 hr and then infiltrated into the lower leaf surface of c. 4 week-old *Ch. quinoa* and spinach (*Spinacia oleracea* cv. Perpetual) plants using a 1ml syringe (without a needle).

Alternatively, the WLV infectious full-length cDNA plasmid constructs were introduced into epidermal cells by biolistic bombardment using a hand gun, essentially as described by Gal On et al. (1997). Thereafter, the plants were grown in a heated glasshouse (18/14°C, 16 hr/8 hr; day/night) for four to seven days before examination.

### Confocal microscopy

Imaging of GFP was performed on a Zeiss LSM 710 upright confocal laser scanning microscope (CLSM; Zeiss Jena, Germany) using a 20x objective, with GFP excitation at 488 nm, emission at 500 to 530 nm.

### Detection and quantitation of WLV by RT-PCR and qRT-PCR

RNA was extracted from leaves of wineberry and bramble plants in the vicinity of JHI and from leaves of *C. quinoa* and spinach plants agroinfiltrated with the WLV cDNA clones using the Machery-Nagel Nucleospin RNA Plant kit. For agroinfiltrated leaves the on-column DNase I step was included, as was a second DNase I treatment of the purified RNA, which was then collected by precipitation with 2.5 M LiCl. RNA extracted from systemic leaves of agroinfiltrated *Ch. quinoa* plants was not DNase I-treated. cDNA was synthesised using the LunaScript kit (New England Biolabs). PCR amplification of WLV was done using PuReTag Ready-to-Go PCR beads (Illustra) or Go Taq DNA polymerase (Promega) according to the manufacturers’ instructions and WLV-specific primer pairs 4194+4195, 4305+4306 or 4361+4264 (Supplementary Table 1). Quantitative PCR was done using the Luna Universal qPCR Master Mix (New England Biolabs) with WLV primers 4361+4264 and as the reference gene spinach Actin 11 (primers 4614+4615; Yu *et al*., 2021), and analysed on a StepOne Real-Time PCR System (Applied Biosystems). Relative levels of WLV RNA accumulation were calculated from triplicate reactions using the ΔΔCt method.

## Results and discussion

### Sequence characterisation of WLV

The ONT sequencing produced 144.7K reads totalling 186.7 Mb. Using a threshold of >80% protein sequence identity one 775 nucleotide (nt) read was identified that aligned to the replication-associated polyprotein (Genbank ID: AEI17897.1) from the allexivirus Blackberry virus E (BVE) (Sabanadzovic *et al*. 2011). All the trimmed reads were then re-aligned to the complete genome of BVE (Genbank: JN053266.1) using Minimap2 (v 2.17) (Li, 2018) which revealed 4 additional reads aligning to that virus. The five reads ranged from 524 to 3257 nt in size and shared between 69 and 85% nucleotide sequence identity with BVE. Using primers derived from these sequences and others developed during the subsequent Sanger sequencing process it was possible to amplify, clone and sequence the entire WLV genome.

This work revealed that the WLV genome (GenBank accession MZ944847) is 7760 nt in size (compared to 7718 nt for BVE) with a 5’ terminal GAAAA sequence that is present also in BVE and some other allexiviruses such as Papaya virus A (PVA) and Shallot virus X (ShVX), and a 3’ terminal polyA tail (Fig. 2). The 5’ untranslated region (UTR) is 97 nt and the 3’ UTR is 111 nt. The first open-reading frame (ORF1) extends from nt 98 to 4636 and encodes a 1513 amino acid (aa) RNA-dependent RNA polymerase protein of 170.6 kDa molecular weight. ORFs 2, 3 and ORFx encode Triple Gene Block (TGB) proteins that are the cell-to-cell movement proteins encoded by a large number of viruses in the *Alphaflexiviridae* and *Betaflexiviridae* families (Verchot-Lubicz *et al*., 2010). ORF2 extends from nt 4667 to 5404 and encodes a 246 aa/27.2 kDa TGB1 RNA binding/RNA helicase protein. ORF3 extends from nt 5385 to 5696 and encodes a 104 aa/11.4 kDa TGB2 protein. ORFx extends from nt 5428 to 5826 and encodes a 133aa/13.9 kDa TGB3-like protein. The TGB1 and TGB2 genes possess AUG (methionine) initiation codons, however, the TGB3-like ORF (denoted as ORFx for BVE; Sabanadzovic *et al*., 2011) overlaps the TGB2 ORF and ORF4 but does not have any AUG codons making it difficult to predict the exact N-terminus of this putative protein. Studies with ShVX suggested that the TGB3-like ORF was expressed by leaky scanning of a subgenomic RNA by ribosomes initiating upstream of the TGB2 gene (Lezzhov *et al*., 2015). This work proposed a CUG (leucine) codon, which was conserved in position in ShVX and five other garlic allexiviruses, as being the translation initiation codon for the TGB3-like protein. This specific leucine is not present in WLV ORFx or the TGB3-like ORFs of four of five other allexiviruses, however, there are clear amino acid sequence similarities in the TGB3-like proteins encoded by all of these viruses. WLV ORF4 extends from nt 5810 to 6877 and encodes a 356 aa/38.9 kDa (P40) protein that is characteristic of all allexiviruses (Supplementary Fig. 1). ORF5 extends from nt 6939 to 7649 and encodes a 237 aa/25.2 kDa coat protein (CP). We used the G-tailing method described above to look for a subgenomic RNA that could function as a messenger for CP expression and obtained an amplicon that began at nt 6896 (5’-GAATCACCAACTAACAAC….) (Fig. 2). This sequence has some similarities with the 5’ terminus of the WLV genomic RNA and is located 53 nt upstream of the CP ORF. WLV, similar to BVE, PVA, Arachis pintoi virus, Alfalfa virus S, Vanilla latent virus and Cassia mild mosaic virus, does not have the ORF for a small nucleic acid-binding protein that is located downstream of the CP gene in allium-infecting allexiviruses. WLV and BVE have CP and replicase protein sequences with, respectively, 99% and 91% identity suggesting WLV is an isolate of BVE (Supplementary Table 2). We designed the primer pair 4194+4195 to amplify a 375 nt region at the 5’ end of ORF1, and primer pair 4035+4036 to amplify a 436 nt region at the 3’ end of the P40 gene and the 5’ end of the CP gene (Supplementary Table 1). These primers detected WLV in the *Ch. quinoa* plants infected using the freeze-dried WLV inoculum. WLV was not detected in any of seven wineberry plants maintained at JHI or in six wineberry plants growing wild in the local area. However, WLV was detected in two wild bramble (*R. fruticosus*) plants growing near to JHI (Fig. 3). The amplified WLV fragments from these plants were cloned and sequenced and found to be more than 99% identical to the freeze-dried virus obtained originally from wineberry.

**Figure 2.**
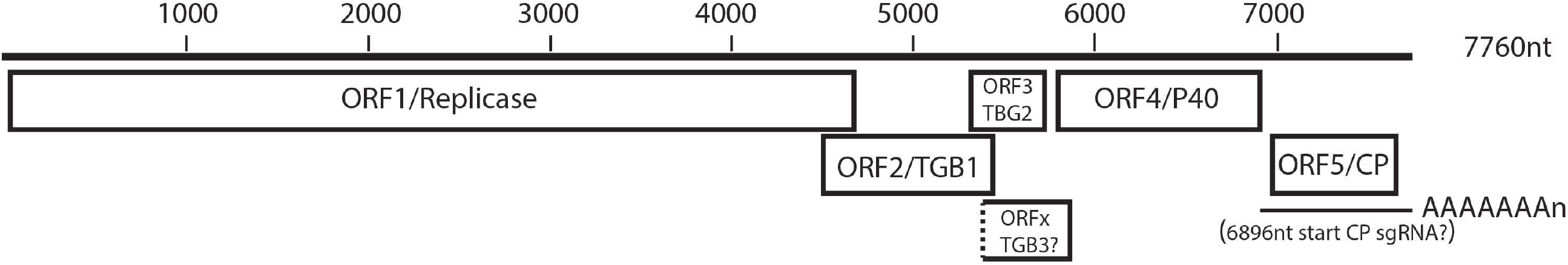
Genome diagram of WLV. ORFs are denoted by rectangular boxes. The N-terminal position of ORFx is uncertain and is shown by a dotted line. The position of the putative subgenomic RNA for CP expression is denoted by a narrow, horizontal line.

**Figure 3.**
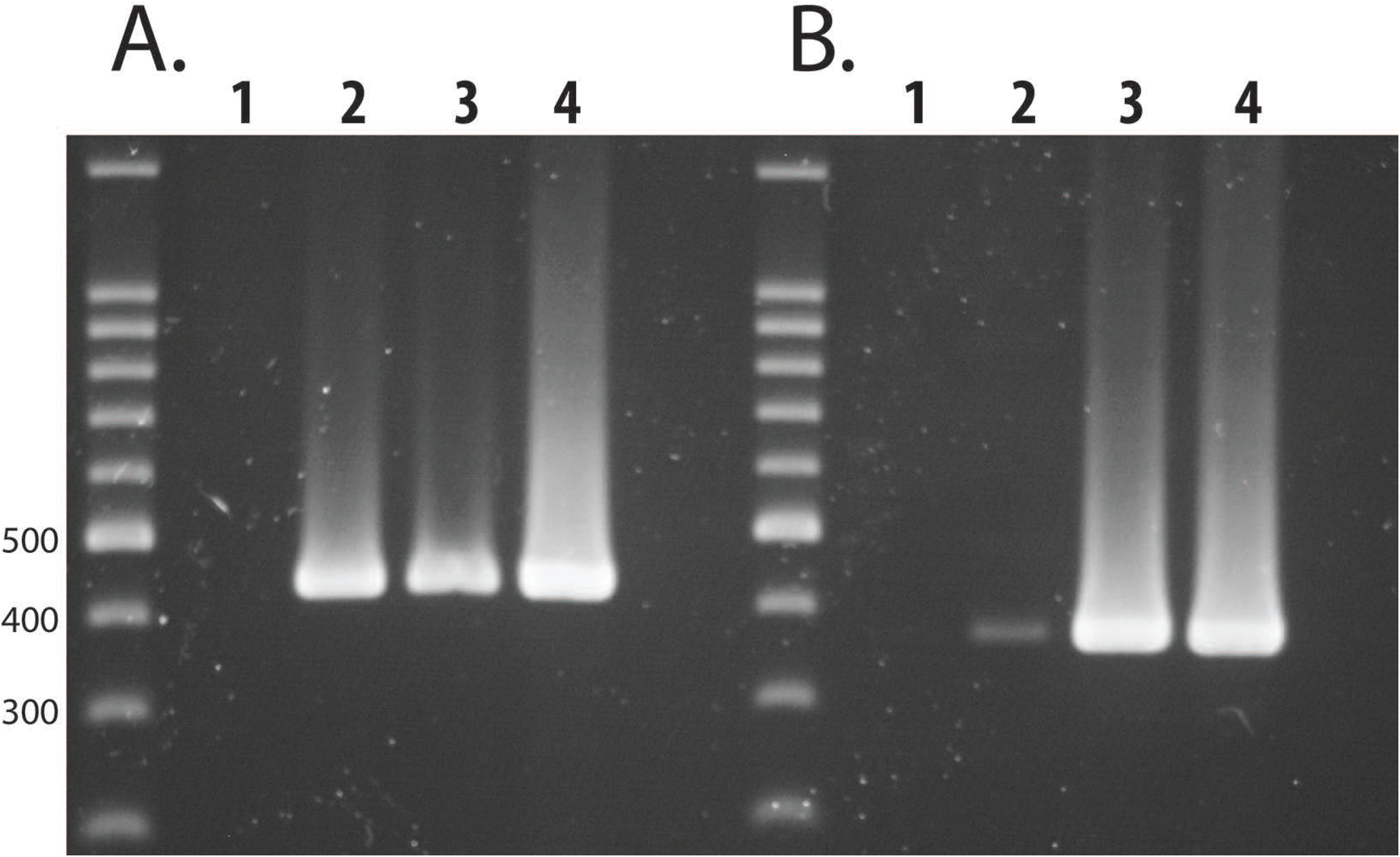
RT-PCR detection of WLV. Lane 1, healthy red raspberry; Lane 2, bramble plant 1; Lane 3, bramble plant 2; Lane 4, *Ch. quinoa* with necrotic lesions. Panel A. primers 4305+4306. Panel B. primers 4194+4195. 100bp DNA markers (Promega).

### Studies with a WLV infectious cDNA clone

Several full-length cDNA clones of WLV were constructed in the small, low-copy vector pDIVA from PCR-amplified virus RNA and a 3’ terminal synthetic 42nt polyA sequence. These clones were infectious in *Ch. quinoa* plants, producing chlorotic patches in agroinfiltrated leaves and sporadic chlorotic spots in systemically-infected leaves (Fig 4., Supplementary Fig. 2). Infection of *Ch. quinoa* was strongly affected by the time of year, with systemic infection occurring only during mid-summer. Agroinfiltration of spinach plants was more reliable, producing a non-symptomatic infection all year round, however, as previously noted WLV did not cause systemic infection in this plant. WLV did not infect *Nicotiana benthamiana or N. clevelandii*.

**Figure 4.**
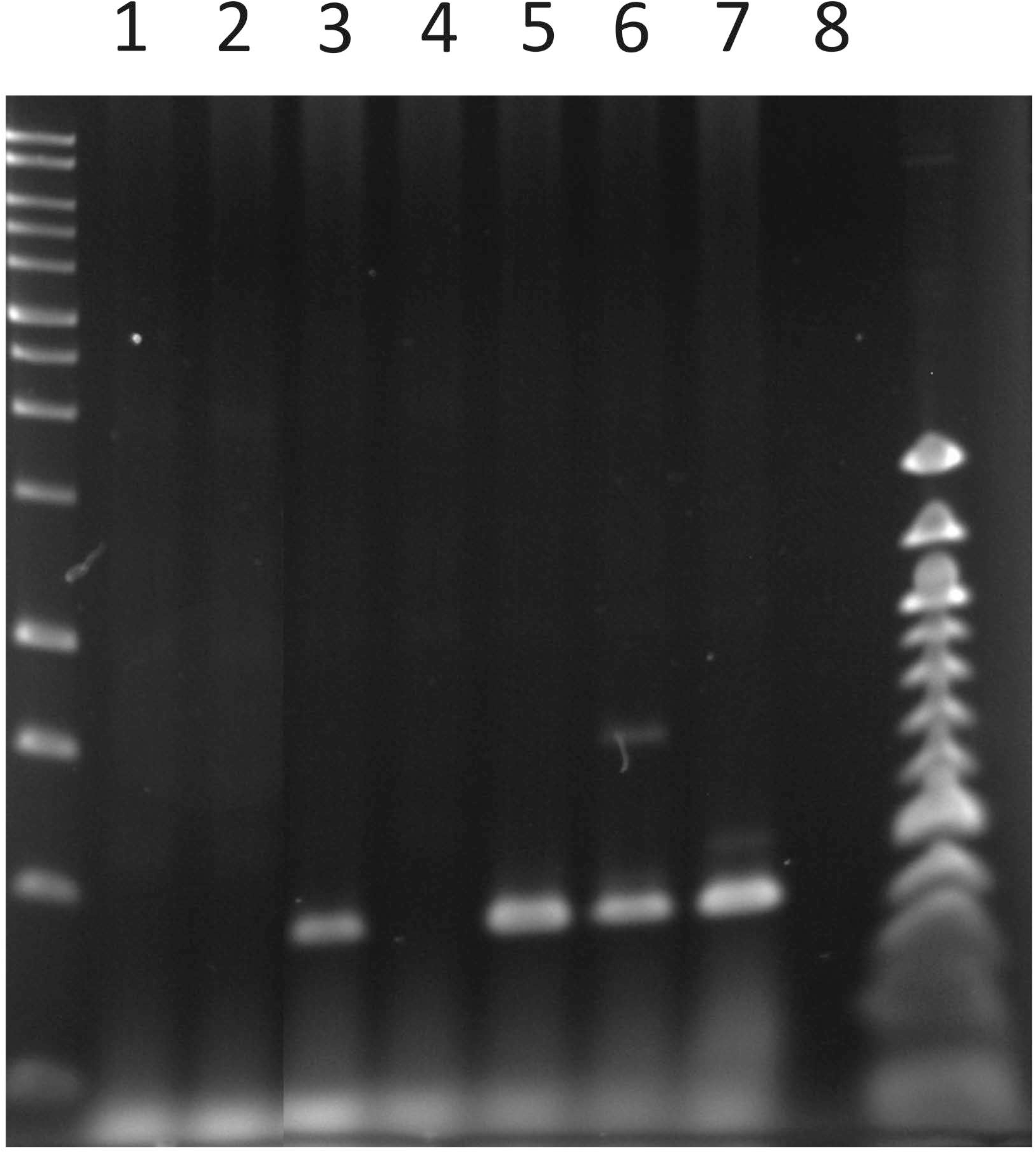
RT-PCR of systemic infection by full-length WLV cDNA clones in *Ch. quinoa* at 12dpi. Lanes 1, 2, healthy plants. Lanes 3, 4, 5, 6, 7 are clones 3.5, 3.6, 5.7, 5.8, 5.9. Lane 8, no RNA. Left lane, 1kb DNA ladder. Right lane, 100 bp ladder.

We introduced frameshift mutations into the WLV ORF1 and P40 genes and examined the accumulation of these mutants and wild-type WLV in infiltrated spinach leaves. PCR amplification of DNase I-treated RNA from these leaves did not produce an amplicon for any of the three infiltrations showing that no residual plasmid DNA from the infiltrated agrobacteria remained in these samples, whereas, WLV was readily amplified from cDNA made from these RNAs (Supplementary Fig. ZZ). Quantitative RT-PCR showed that the P40 mutation reduced the level of WLV RNA in these infiltrated leaves by 97% (Fig. 5). An extremely small amount of WLV RNA (0.25% of wild-type WLV) was detected for the ORF1 frameshift mutant, which we assume represents non-replicating transcripts expressed from the binary plasmid in the infiltrated leaves.

**Figure 5.**
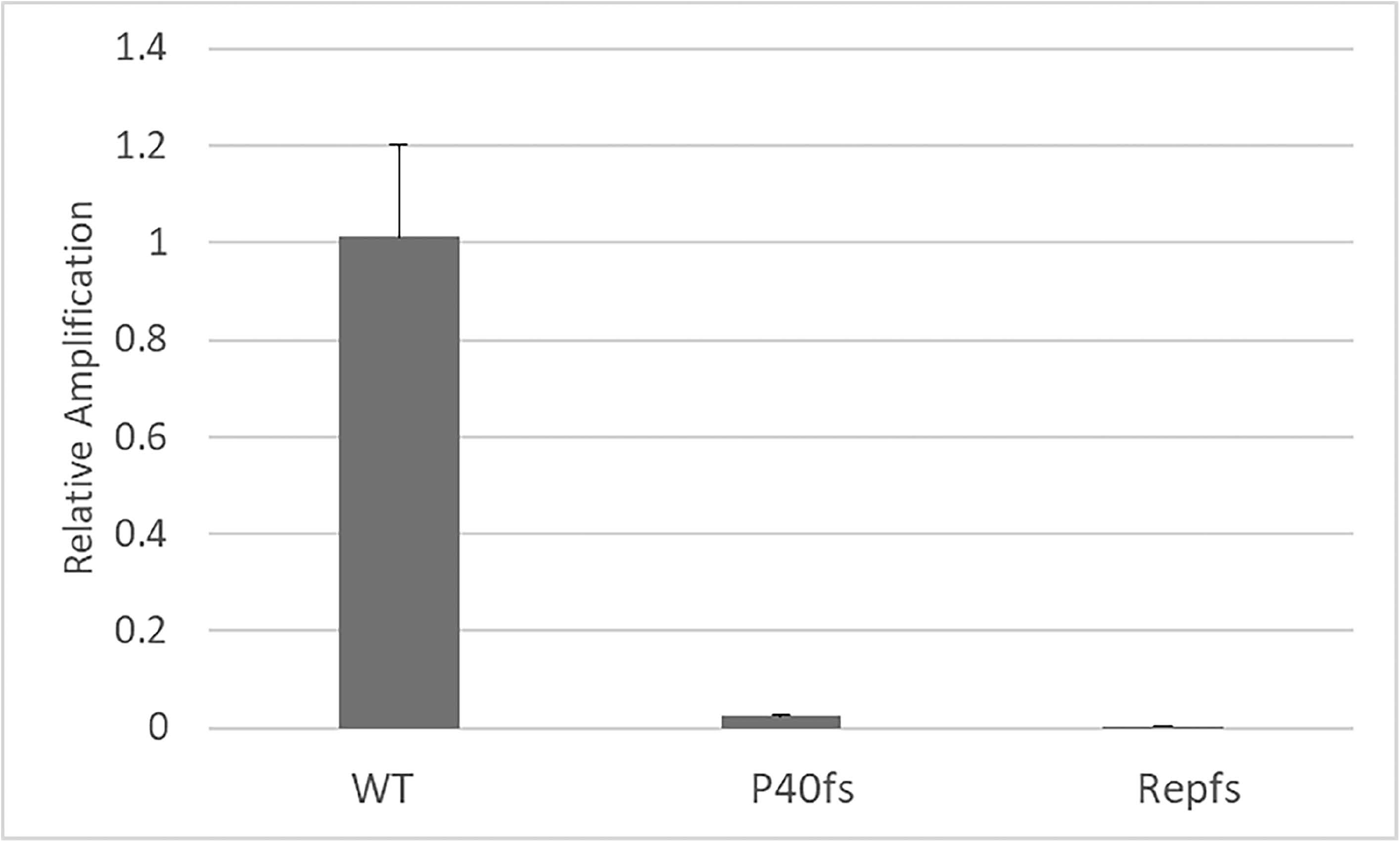
P40 frameshift mutation reduces WLV RNA accumulation by 97% at 7dpi in agroinfiltrated spinach leaves. Error bars show standard deviation.

We constructed a version of WLV in which the CP was expressed with GFP fused at its N-terminus, with the two proteins separated by a self-cleaving FMDV 2A peptide. The 2A cleavage is not 100% efficient and with a different virus, PVX, this approach allowed the formation of (overcoated) virus particles that contained GFP-CP fusion protein interspersed with fully cleaved CP subunits (Santa Cruz *et al*., 1996). This modified PVX was able to move systemically in *N. benthamiana* and *N. tabacum* plants. For the GFP-overcoated WLV the virus was able to multiply and move from cell-to-cell in infiltrated leaves, forming large assemblies of fluorescing cells, but did not move into leaf veins and was never observed in systemic leaves (Fig. 6). This was the case in spinach, where WLV is confined to inoculated leaves, and also in *Ch. quinoa*, where WLV is known to occasionally cause systemic infection. The P40 mutant version of the GFP-overcoated WLV was observed in only a small number of cells in the infiltrated leaves and with a low level of fluorescence, corroborating the very low level of accumulation of the non-overcoated P40fs WLV mutant.

**Figure 6.**
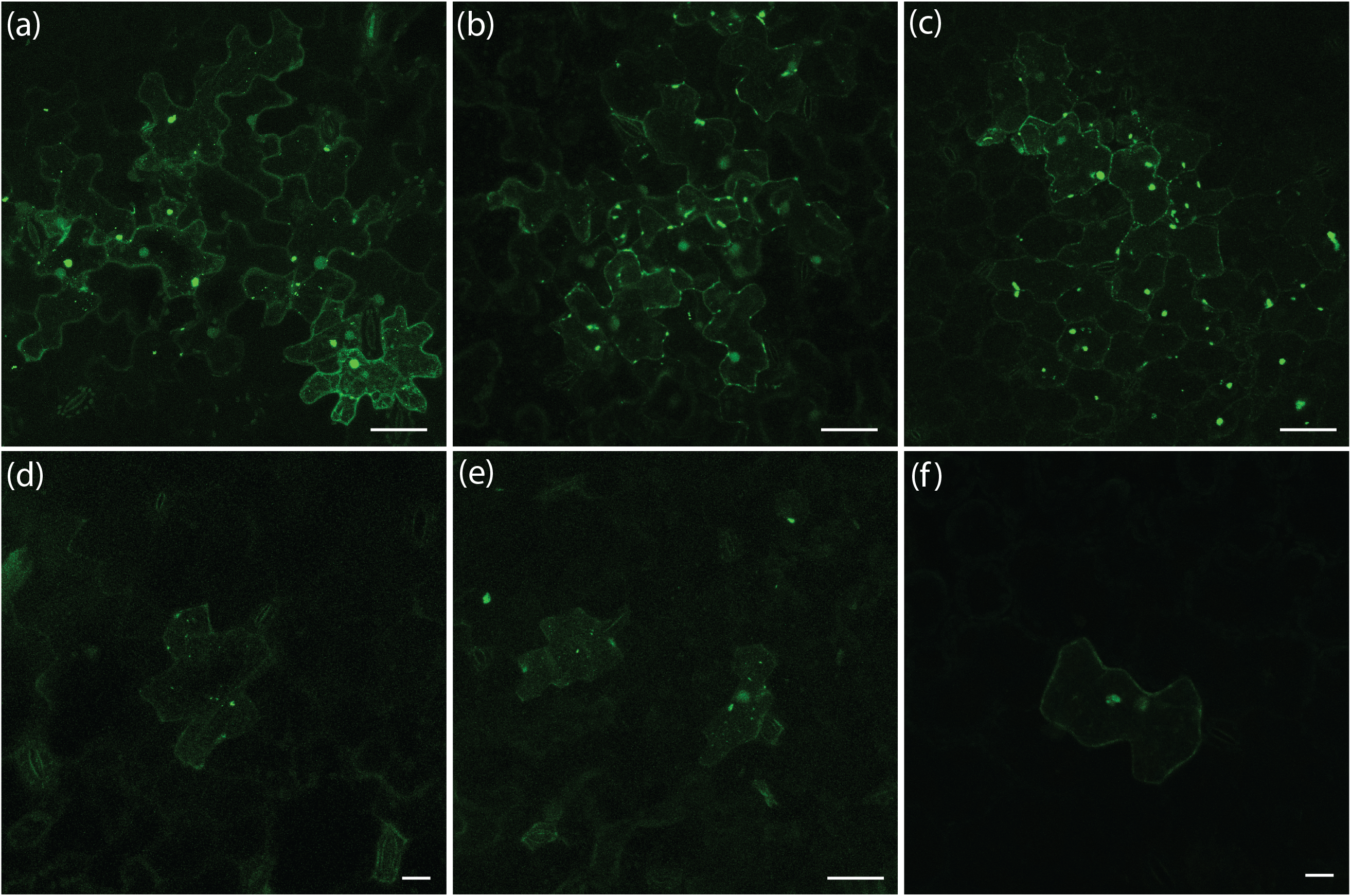
Confocal microscopy images of fluorescence from GFP overcoated WLV. Panels a to c, wild-type WLV (p2108). Panels d to f, P40fs mutant (p2111). Panel a, agroinfiltrated spinach, 4dpi. Panel b, agroinfiltrated *Ch quinoa*, 7dpi. Panel c, bombarded *Ch. quinoa*, 7dpi. Panels d and e, agroinfiltrated *Ch. quinoa*, 7dpi. Panel f, bombarded *Ch. quinoa*, 7dpi. Size bars are 40 µM (panels a, b, c, e) or 20 µM (panels d, f).

## Conclusion

In this work we have identified and characterised WLV, showing it to be an isolate of the allexivirus BVE. This has enabled us to design several diagnostic RT-PCR primers that can now be used in certification of WLV-free raspberry and blackberry plants for soft fruit breeders and propagators. Although we did not find this virus in wineberry plants in the vicinity of JHI, we did find the virus in local wild bramble plants. The vector for WLV is not known, although other allexiviruses are known or suspected to be transmitted between plants by eriophyid mites. Another raspberry-infecting virus, Raspberry leaf blotch virus, is also transmitted by an eriophyid mite, *Phyllocoptes gracilis*, and causes significant damage to raspberry production (McGavin et al., 2012).

Apart from genome sequencing there have not been very many molecular studies done on allexiviruses. These viruses are divided into two groups depending on their genome organization. The *Allium*-infecting allexiviruses have genomes encoding ORF1 (putative replicase), ORF2 (TGB1), ORF3 (TGB2), ORFx (a potential non-AUG initiated gene), ORF4 (P40), ORF5 (CP) and ORF6 (cysteine rich protein; CRP). WLV, and other allexiviruses including Papaya virus A, Alfalfa virus S and Arachis pintoi virus, encode similar ORFs 1 to 5 but do not encode the ORF6 (CRP). Using a recently described infectious cDNA clone of Garlic virus C both the CRP and CP were shown to act as competing suppressors of RNA silencing (Kim *et al*., 2023). The only previously documented study of the allexivirus-specific P40 protein in WLV) used an infectious clone of Shallot virus X, a CRP-encoding allexivirus, in sugar beet protoplasts (Vishnishenko *et al*., 2000; 2002). A frameshift mutation in the P42 gene did not affect CP production, as shown by western blotting, but abolished the formation of virus particles, however, in these experiments the accumulation of virus RNA was not monitored. Our study of WLV shows that mutation of the P40 gene reduces the accumulation of virus RNA by 97%, although we do not know what effects there are on the level of expressed CP or formation of virus particles. It is possible that, as this group of allexiviruses do not encode the CRP RNA silencing suppressor protein, the WLV P40 might have been adapted to this role, which might explain the extreme effect of the P40 mutation on WLV accumulation. Nevertheless, further studies are required to better understand the function of the WLV P40 protein in the virus life cycle.

## Acknowledgements

We thank Jenny Morris and Pete Hedley for the nanopore sequencing of WLV.

We thank Mark McKillen for help with the WLV cloning.

## Author contributions

SMF devised the project and supervised the work. WM did laboratory work for virus isolation, RT-PCR amplification, cloning and Sanger sequencing. SJ devised and conducted the bioinformatics analysis of the RNA-seq data. GC did the confocal microscopy. SMF wrote the manuscript.

## Figure Legends

**Supplementary Figure 1.**
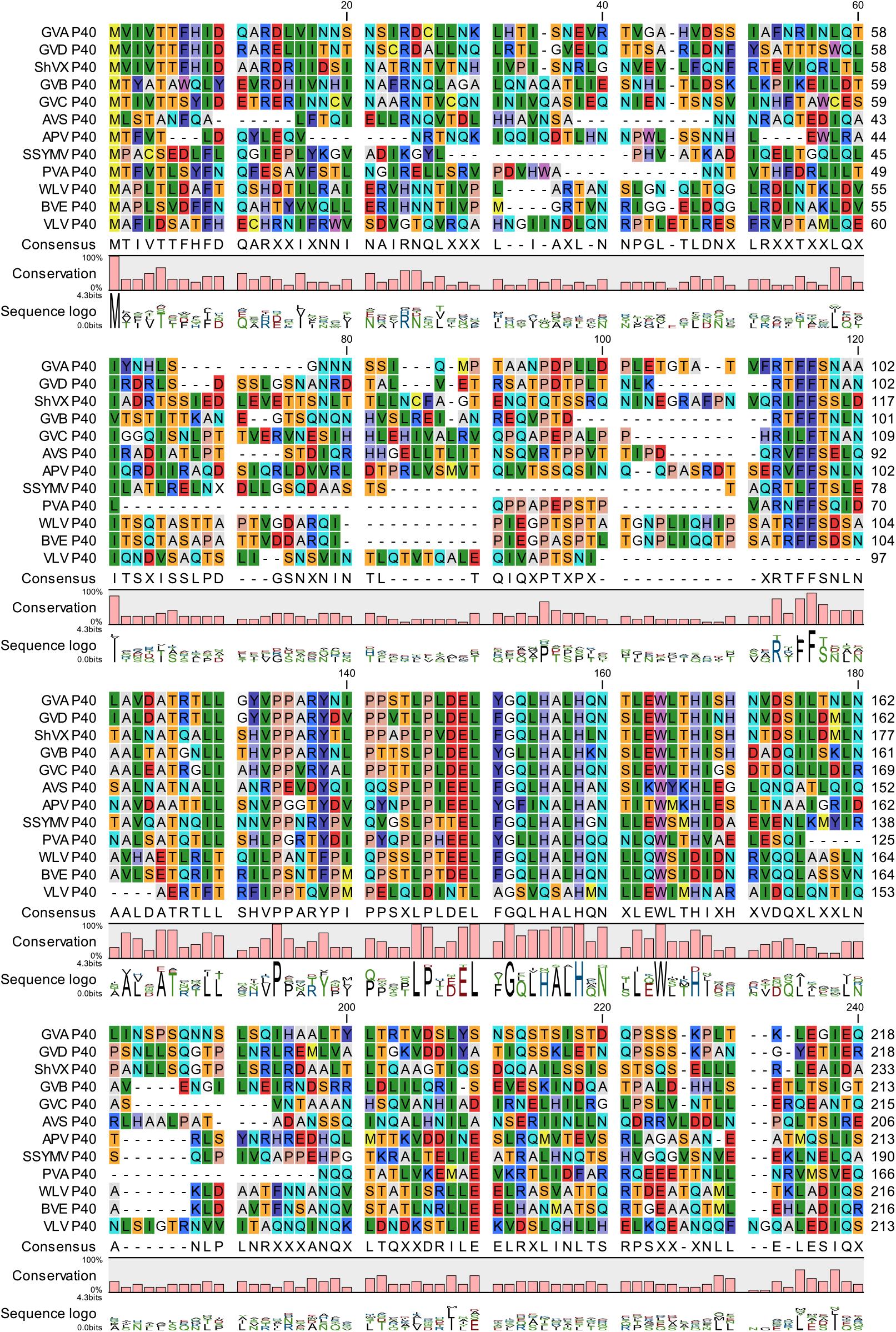
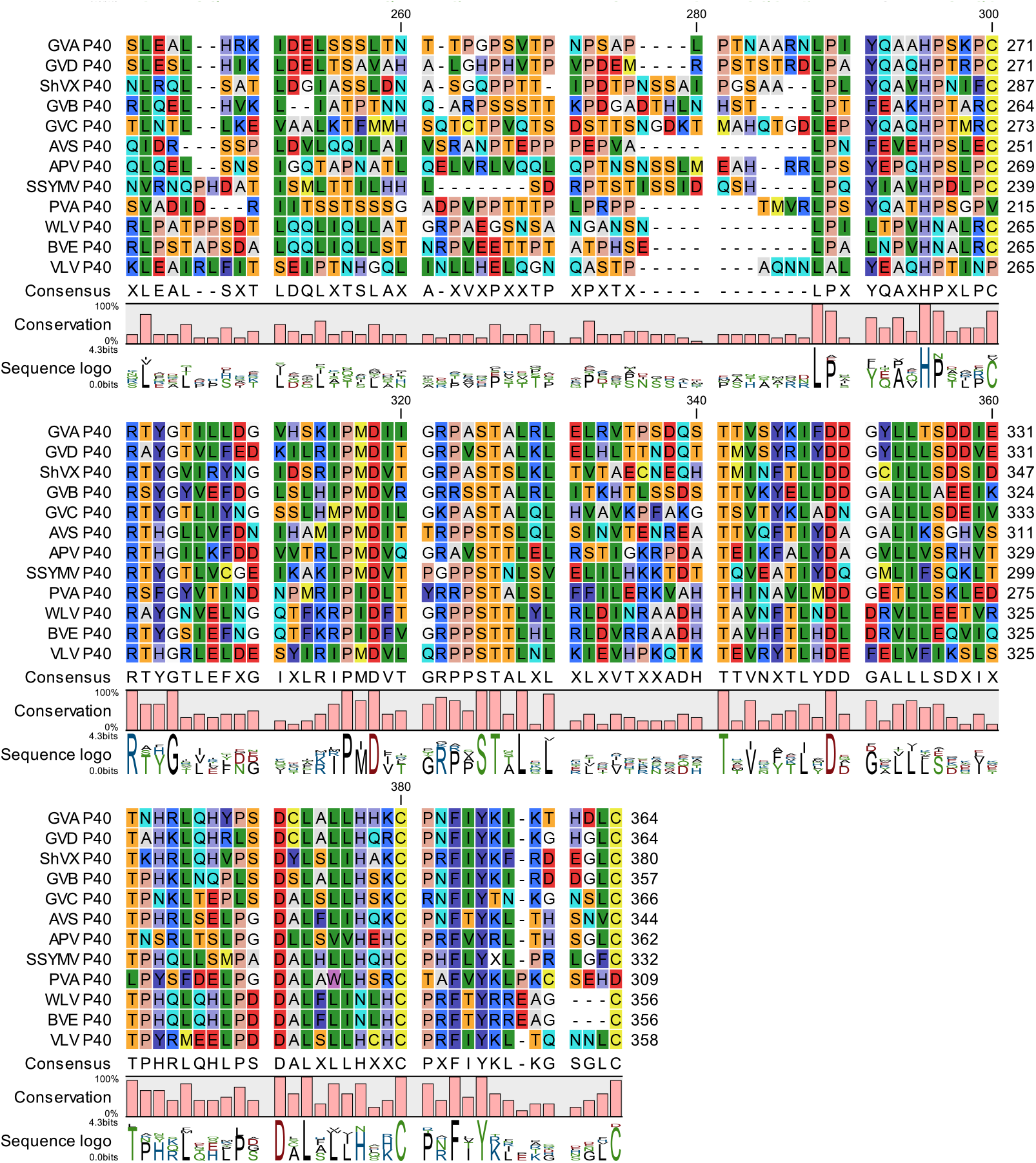
Alignment of P40 proteins encoded by allexiviruses. GVA; Garlic Virus A, QNJ60322.1. GVB; Garlic Virus B,. GVC; Garlic Virus B, YP_009110671.1. GVC; Garlic Virus C, UUQ75122.1. GVD; Garlic Virus D, QXE32444.1. ShVX; Shallot Virus X, UTU35108.1. AVS; Alfalfa Virus S, QJD13461.1. APV; Arachis Pintoi Virus, YP_009328895.1. SSYMV; Senna Severe Yellow Mosaic Virus, QEM20972.1. PVA; Papaya Virus A, QIM41189.1. WLV; Wineberry Latent Virus, UYK33002.1. BVE; Blackberry Virus E, YP_004659203.1. VLV; Vanilla Latent Virus, ASJ78781.1.

**Supplementary Figure 2.**
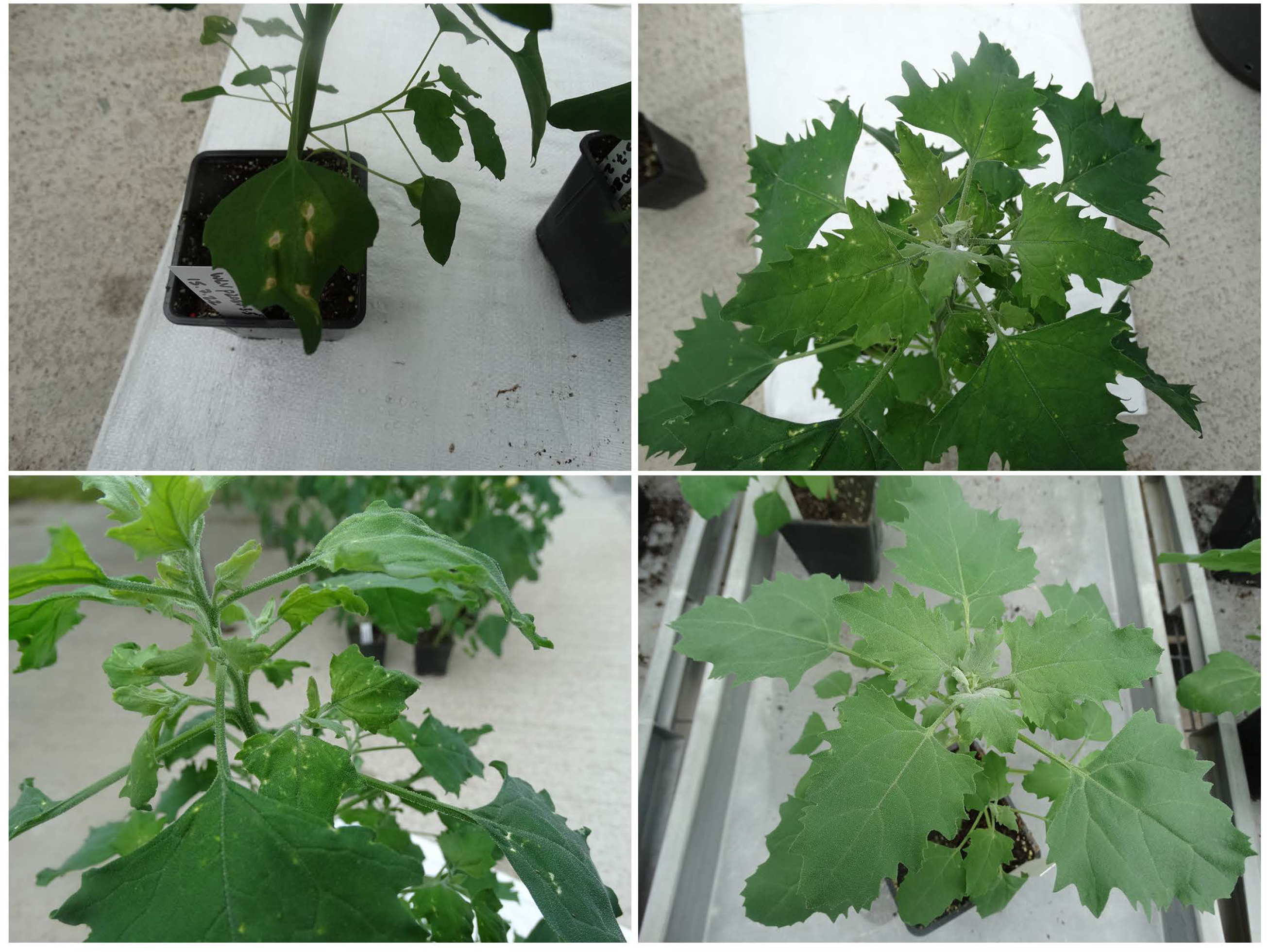
Symptoms at 12 dpi on *Ch. quinoa* plants after infection using agrobacterium-delivered full-length cDNA clones of WLV. Top left, infiltrated leaf. Top right and lower left, systemic chlorotic spots. Lower right, uninoculated plant.

## Primers used in this work

Oxford Nanopore sequencing

PCR-RH-RT; TTTCTGTTGGTGCTGATATTGCTGCCATTACGGCC[mG][mG][mG]

PCR-Sw_mod_3G; AAGTTCATTTCATTTGGAGAGGACAAAATATAAGAACGAAGTATTATGTTCG

Construction of full-length clone

4347; AAGTTCATTTCATTTGGAGAGGGTTTCAGGGTTCTACGTAACCCCC

4104; GAGATGCCATGCCGACCC

4344;

GTTTCAGGGTTCTACGTAACCCCCAAAAAAAAAAAAAAAAAAAAAAAAAAAAAAAAAAAAAAAAAAAAAAAA AGGGTCGGCATGGCATCTCCACCTCCTCGCG

2740; CCTCTCCAAATGAAATGAACTTCC

2757; GGGTCGGCATGGCATCTCCA

Construction of GFP:CP overcoat

4456; CTCCATTTACCATCTGTTCATACACCATGAGTAAAGGAGAAGAACTTTTCACTGGAG

4458; GGAGGGTTAGCTGGTGTGTCAGGCCCAGGGTTGGACTCGACGTCTC

WLV qRT-PCR quantitation/detection

4194; AGTGCGAAGACTTCTCGACC

4195; ATCCCGTACCGCAAGACGT

4305; CTTTACCTCCGCCTGGACAT

4306; GGTGGCTACCTTGTTGGCT

4264; TACATACCACACGGTGAGGG

4361; GTCGATAGGCCCCACATCTG

Spinach Actin 11 qRT-PCR quantitation

4614; CGAGCTGTGTTCCCTAGTATTG

4615; CGAGCTGTGTTCCCTAGTATTG

---

**Supplementary Table 2.**
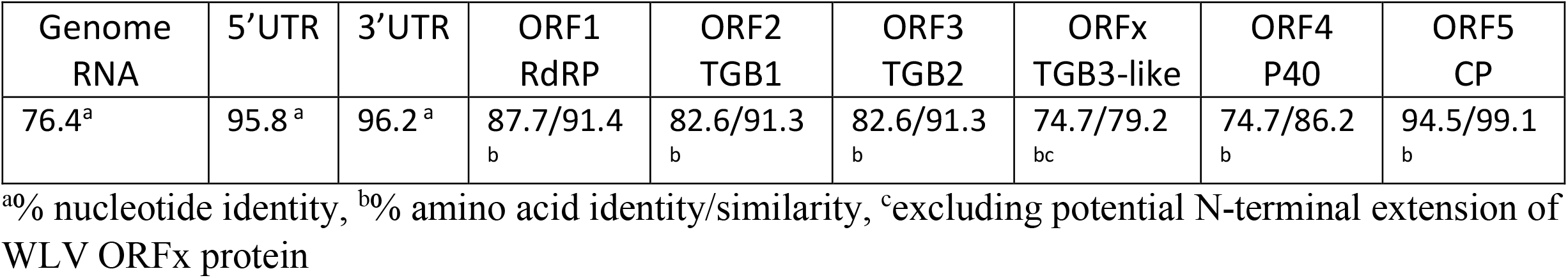
Nucleotide and amino acid identities between WLV and BVE RNAs and predicted proteins.

| Genome RNA | 5'UTR | 3'UTR | ORF1 RdRP | ORF2 TGB1 | ORF3 TGB2 | ORFx TGB3-like | ORF4 P40 | ORF5 CP |
| --- | --- | --- | --- | --- | --- | --- | --- | --- |
| 76.4<sup>a</sup> | 95.8<sup>a</sup> | 96.2<sup>a</sup> | 87.7/91.4<sub>b</sub> | 82.6/91.3<sub>b</sub> | 82.6/91.3<sub>b</sub> | 74.7/79.2<sub>bc</sub> | 74.7/86.2<sub>b</sub> | 94.5/99.1<sub>b</sub> |

<sup>a</sup>% nucleotide identity, <sup>b</sup>% amino acid identity/similarity, <sup>c</sup>excluding potential N-terminal extension of WLV ORFx protein

## References

De Coster, W., D’Hert, S., Schultz, D. T., Cruts, M. & Van Broeckhoven, C. NanoPack: Visualizing and processing long-read sequencing data. Bioinformatics 34, 2666–2669 (2018). DOI: 10.1093/bioinformatics/bty149

Gal On, A., Meiri, E., Elman, C., Gray, D. J., and Gaba, V. 1997. Simple hand-held devices for the efficient infection of plants with viral-encoding constructs by particle bombardment. J. Virol. Methods 64:103–110. 10.1016/S0166-0934(96)02146-5

Jones, A.T. 1977. Partial purification and some properties of wineberry latent, a virus obtained from Rubus phoenicolasius. Ann. Appl. Biol. 86, 199–208. 10.1111/j.1744-7348.1977.tb01832.x

Jones, A.T., Mitchell, M.J., McGavin, W.J., Roberts, I.M. 1990. Further properties of wineberry latent virus and evidence for its possible involvement in calico disease. Ann. Appl. Biol. 117, 571–581. 10.1111/j.1744-7348.1990.tb04823.x

Kim H, Kawakubo S, Takahashi H, Masuta C (2023) Two mutually exclusive evolutionary scenarios for allexiviruses that overcome host RNA silencing and autophagy by regulating viral CRP expression. PLoS Pathog 19(6): e1011457. 10.1371/journal.ppat.1011457

Lanfear, R., Schalamun, M., Kainer, D., Wang, W. & Schwessinger, B. MinIONQC: Fast and simple quality control for MinION sequencing data. Bioinformatics 35, 523–525 (2019). DOI: 10.1093/bioinformatics/bty654

Laufer, M., Mohammad, H., Maiss, E., Richert-Pöggeler, K., Dall’Ara, M., Ratti, C., Gilmer, D., Liebe, S., Varrelmann, M. 2018. Biological properties of Beet soil-borne mosaic virus and Beet necrotic yellow vein virus cDNA clones produced by isothermal in vitro recombination: Insights for reassortant appearance. Virology 518, 25–33. 10.1016/j.virol.2018.01.029

Lezzhov, A.A., Gushchin, V.A., Lazareva, E.A., Vishnichenko, V.K., Morozov, S.Y., Solovyev, A.G. 2015. Translation of the Shallot virus X TGB3 gene depends on non-AUG initiation and leaky scanning. J. Gen. Virol. 96, 3159–3164. 10.1099/jgv.0.000248

Li, H. Minimap2: Pairwise alignment for nucleotide sequences. Bioinformatics 34, 3094–3100 (2018). DOI: 10.1093/bioinformatics/bty191

McGavin, W.J., MacFarlane, S.A. 2009. Rubus chlorotic mottle virus, a new sobemovirus infecting raspberry and bramble. Vir. Res. 139, 10–13. 10.1016/j.virusres.2008.09.004

McGavin, W.J., Mitchell, C., Cock, P.J.A., Wright, K.M., MacFarlane, S.A. 2012. Raspberry leaf blotch virus, a putative new member of the genus Emaravirus, encodes a novel genomic RNA. Journal of General Virology, 93, 430 – 437. 10.1099/vir.0.037937-0

Sabanadzovic, S, Abou Ghanme-Sabanadzovic, N., Tzanetakis, I.E. 2011. Blackberry virus E: an unusual flexivirus. Arch. Virol. 156, 1665–1669. DOI: 10.1007/s00705-011-1015-y

Santa Cruz, S., Chapman, S., Roberts, A.G., Roberts, I.M., Prior, D.A.M., Oparka, K.J. 1996. Assembly and movement of a plant virus carrying a green fluorescent protein overcoat. Proc. Natl. Acad. Sci. USA 93, 6286–6290. 10.1104/pp.103.037507

Yu, H., Ma, Y., Lu, Y., Yue, J., Ming, R. Expression profiling of the Dof gene family under abiotic stresses in spinach. Sci. Rep. 11, 14429 (2021). DOI: 10.1038/s41598-021-93383-6

Verchot-Lubicz, J., Torrance, L., Solovyev, A.G., Morozov, S.Y. Jackson, A.O., Gilmer, D. 2010. Varied movement strategies employed by triple gene block-encoding viruses. MPMI 23, 1231–1247. DOI: 10.1094/MPMI-04-10-0086

Vishnichenko, V.K., Kaloshin, A.A., Ryabov, E.V., Zavriev, S.K. 2000. Cloning of full-length cDNA of the Shallot virus X genome and infectivity of its transcripts in sugar beet protoplasts. Molecular Biology, 34, 148–151.

Vishnichenko, V.K., Stel’mashchuk, V.Y., Zavriev, S.K. 2002. The 42K protein of Shallot virus X participates in formation of virus particles. Molecular Biology, 36, 879–882.

Young, K.T., Lahmers, K.K., Sellers, H.S., Stallknecht, D.E., Poulson, R.L., Saliki, J.T., Tompkins, S.M., Padykula, I., Siepker, C., Howerth, E.W., Todd, M., Stanton. J.B. 2019. Randomly primed, strand-switching MinION-based sequencing for the detection and characterization of cultured RNA viruses. bioRxiv 2019.12.16.875872; doi: 10.1101/2019.12.16.875872

